# The cuticle lamellae responsible for structural coloration in red algae display common features with the metazoan extracellular matrix

**DOI:** 10.64898/2026.01.12.698930

**Authors:** Glenn Philippe, Olivier Godfroy, Fred Beisson, Ambre Gautier, Diane Jouanneau, Sophie Le Panse, Emmanuelle Com, Ludovic Delage, Mirjam Czjzek, Elizabeth Ficko-Blean, Jonas Collén

**Affiliations:** Sorbonne Université, CNRS, Laboratoire de Biologie Intégrative des Modèles Marins, LBI2M, UMR8227, Station Biologique de Roscoff, Place Georges Teissier F-29680 Roscoff, Bretagne, France; CEA, CNRS, Aix Marseille Université, Institut de Biosciences et Biotechnologies d’Aix-Marseille, BIAM, UMR7265, CEA Cadarache, Saint-Paul-lez-Durance F-13108, France; CNRS, FR 2424, Merimage Facility, Station Biologique de Roscoff, Place Georges Teissier F-29680 Roscoff, Bretagne, France; Univ Rennes, CNRS, Inserm, Biosit UAR 3480, US-S 018, Protim Core Facility, Rennes 35042, France

## Abstract

Structural coloration is a physical phenomenon observed in many living organisms, including seaweeds, via the formation of highly organized nanostructures. Although the deposition of multilayer thin-film in the extracellular matrix (ECM) is a widespread characterized system, enabling light interference, to date such lamellae have only been described in the cuticle of red algae belonging to the Gigartinaceae family within macroalgae. Here, we determine the composition of the multilayered cuticle components in the model species *Chondrus crispus* (Rhodophyta), using genomic, transcriptomic, proteomic, metabolomic and analytical profiling of carbohydrates. The complex structural assembly revealed common features with the ECM of animals. The carbohydrate fraction consists of a complex mixture of ECM carrageenans and glycosaminoglycan-like structures. A major ‘von Willerbrand factor A’ domain (VWA) protein plays a critical role for the adhesion of macromolecules, including protein-protein interactions and to sulfated polysaccharides. We propose the name of Lamellae Cohesive Protein (LCP) regarding the alternate distribution of chemically distinct constituents within the cuticle. For the first time, we identified major proteins of the algal cuticle, providing a framework for addressing the evolutionary origins of the algal cuticle biosynthesis and raise important questions regarding its role, particularly across the life cycle marked by major structural ECM modifications.

## Introduction

Structural coloration produced by lamellae in the epidermal ECM has been described in multicellular organisms across multiple phyla, including plants and animals, where it results from the reflection of specific wavelengths that are enhanced by a physical mechanism of constructive interference (Parker and Martini 2006; Vignolini et al. 2013). These multilayer reflectors alternate in refractive index through the arrangement of chemically distinct components that each organism sources from its own epidermal extracellular matrix (ECM), for example, see (Seago et al. 2009; Dobson et al. 2025; Middleton and Sinnott-Armstrong 2024). Few studies have investigated structural color in macroalgae; consequently, little is known about chemical composition, structural mechanisms, and biological functions of their multilayer reflectors.

The algal cuticle is a constitutive protein-rich layer covering the surface of some red, green and brown macroalgae (Hanic and Craigie 1969). It is formed on the outermost part of the ECM, creating a resistant boundary with the environment. Considerable variation is known in the composition and biosynthesis of the ECM of macroalgae, which arise from their distinct evolutionary histories (Kloareg et al. 2021), however, the biochemistry and structure of the algal cuticle is scarcely documented. Unknown proteinaceous material has been shown to be predominant in the resistant structure of all algal cuticle, and may account for up to 80% of the composition (Hanic and Craigie 1969). ECM polysaccharides have been reported as important cuticle components in some red algal species (Frei and Preston 1964; Gerwick and Lang 1977; Craigie et al. 1992; Flores et al. 1997). The analogous plant cuticle of the Embryophytes is composed of the insoluble lipid polyester cutin and hydrophobic waxes that prevent desiccation and facilitate terrestrial life (Yeats and Rose 2013); however, neither structure nor primary function are shared with algal cuticles.

In the Rhodophyta, a multi-layered cuticle that is responsible for structural coloration has been reported as part of an algal-specific photoprotective mechanism a single family of red seaweeds, the Gigartinaceae (Chandler et al. 2017; Fleitas et al. 2024; Arnould-Pétré et al. 2025). The ECM of Gigartinaceae is distinguished by containing sulfated galactans, carrageenans, which present important molecular structure variations between phases of the red algal haplodiplontic life cycle (McCandless et al. 1983; van de Velde 2008). Although significant structural variation in carrageenans leads to distinct rheological properties, no effects on thallus morphology have been characterized between gametophytes (haploid, 1n) and tetrasporophyte (diploid, 2n) individuals, including in the isomorphic model species *Chondrus crispus*, where different carrageenans are predominant in the gametophytes and tetrasporophyte ECM (Chopin and Floc’h 1992; García Tasende et al. 2011; Ropartz et al. 2025). Fundamental to this study, the ultrastructure of the cuticle lamellae also varies throughout *C. crispus* life cycle, thereby producing a structural coloration exclusive to the gametophyte (Craigie et al. 1992; Chandler et al. 2015). Notably, Craigie et al. (1992) reported differences in the protein content of the cuticle between the gameophyte and tetrasporophyte stages, which likely participate in the variation of the multi-layered cuticular arrangement. This hypothesis implies that functions inherent to the cuticle is mediated by the assembly of specific compounds, which would have particularly specific assembly properties at the interface with the environment.

In the present study, we aimed to decipher the role of cuticular components in the algal cuticle formation of *C. crispus.* After optimizing the rapid isolation of an intact cuticle, a series of analyses were undertaken to explore its biochemical nature, which uncovered a novel algal glycosaminoglycan-like. Using a proteomic approach, we identified a major structural protein specific to the *C. crispus* gametophyte individuals. This protein consists of a unique von Willebrand factor A domain (VWA) that is widespread in all Eukaryotes and commonly found in various proteins of the Metazoan ECM (Whittaker and Hynes 2002). As a candidate to the cuticle assembly, we investigate properties of this protein to form proteinaceous layers and studied its interactions with polysaccharides. Our data highlight a key role of this protein in the specific architecture of algal cuticles that produce structural coloration, and offers insights into a conserved function in the cuticle assembly beyond the Gigartinaceae species.

## Results

### Rapid isolation of *Chondrus crispus* cuticle preserves organized lamellae

The blue coloration of *C. crispus* occurs mainly on the tip of the thallus of gametophyte individuals, where the ultrastructure of the cuticle shows greater number and parallel electron-dense lamellae (Fig. 1a, b, c) (Chandler et al. 2015). The optimization of a protocol to chemically isolate the multi-layered cuticle aimed to overcome the constraints associated with mechanical peeling. Using a rapid and high-yield protocol, based on incubation in an adjusted concentration of sulfuric acid, structural integrity of the cuticle lamellae of *C. crispus* is preserved, as evidenced by the blue reflections of the isolated cuticle, as well as inter-lamellae fibrillar material that is a feature of the fine ultrastructure (Gerwick and Lang 1977; Craigie et al. 1992) (Fig. 1d, e, f). Moreover, vesicle-like structures, which are observed throughout the epidermal ECM and seem to accumulate at the cuticle level, are still noticeable on the isolated cuticle (Fig. 1c, e). The algal cuticle is reported to be generally highly resistant to strong acids (Hanic and Craigie 1969), however, the lamellae organization of *C. crispus* cuticle is highly altered when submitted to excessive concentration, which results in the loss of structural coloration (Fig 1g, Supplementary Fig. 1).

**Figure 1.**
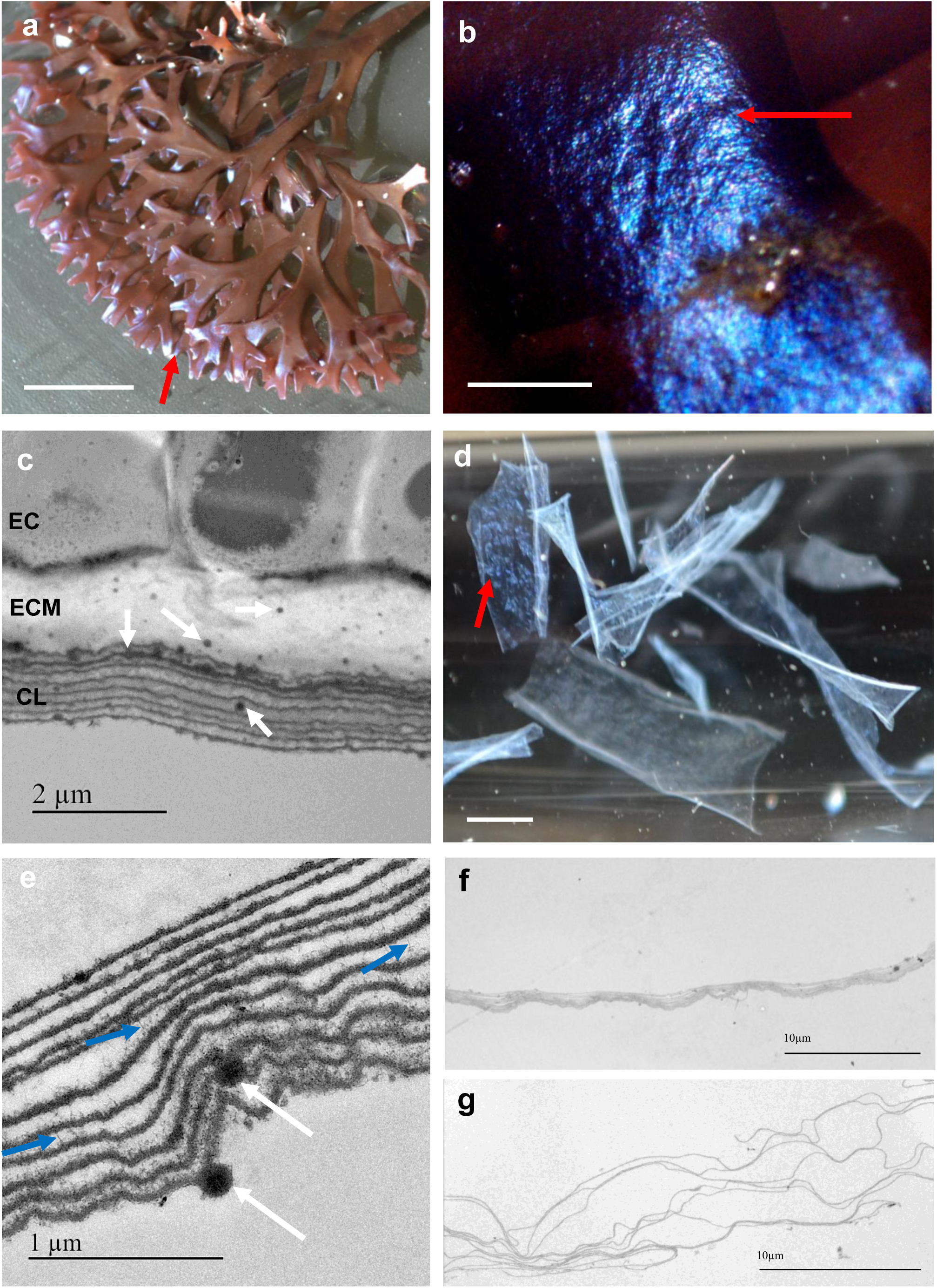
Isolation of cuticle lamellae. **a** Gametophyte thallus of the red algal model *Chondrus crispus* exhibiting structural coloration on tips (red arrow). Scale bar = 20 mm. **b** Zoom on the typical metallic blue coloration produced at the surface of the tips. Scale bar = 1 mm. **c** TEM cross-section of the epidermis showing epidermal cells (**EC**) extracellular matrix (**ECM**), and the multilayered cuticle (**MC**) responsible for coloration by light interference. White arrows highlight vesicle-like structures throughout the ECM and CL. Scale bar = 2 µm. **d** Cuticle films in suspension after chemical isolation, displaying structural coloration. Scale bar = 2 mm. **e, f, g** TEM cross-sections of cuticle after chemical isolation. **e, f** Preserved lamellar organization following optimized conditions of isolation (35% H2SO4). **e** Blue arrows highlight narrow filaments connecting electron-dense layers. White arrows indicate vesicles incorporated into the assembly. Scale bar = 1µm. **f** Large view of the material with scale bar = 10 µm. **g** Large view of the material after isolation with excessive acid concentration (70% H2SO4), showing disorganized electron-dense layers. Scale bar = 10 µm.

When applied to tetrasporophytes, the protocol produced varying success rates, with cuticle detachment limited to the far extremity of tips resulting in thread-like pieces (Supplementary Fig. 2). The challenging isolation of small amounts of cuticle from tetrasporophytes is consistent with previous description of layers that anastomose freely, resulting in a flaking assembly, overall less robust and thinner than in the gametophyte (Craigie et al. 1992).

### Proteomic analysis of *C. crispus* cuticle highlights structural candidates for protein-protein interaction

Proteins were extracted from the isolated cuticle using denaturing reagents and separated by SDS-PAGE to generate a protein profile (Fig. 2a). Three major bands in the gametophyte sample ranged between 10kDa and 25kDa, and a major band in the tetrasporophyte sample resolved poorly around 30kDa, which corroborate earlier observations from Craigie et al. (1992). We identified specific sets of proteins for each life cycle stage by LC-MS/MS analysis of trypsin-digested protein extracts (Fig. 2b). Most of the proteins detected were predicted to be secreted proteins based on the presence of a putative signal peptide and using the program DeepLoc-2.1 (Supplementary Table 1). Identified in the tetrasporophyte cuticle are two uncharacterized protein homologs (UniProt accession: R7Q339 and 57Q297), which correspond in their molecular weight to the major band, both proteins have highly specific expression in this stage of the life cycle, with log_2_ fold changes of 9.5 and 9.9 (Lipinska et al. 2020). These proteins appear to be restricted to the Gigartinaceae as no homologs can be detected in any other family to date, thus representing a lineage-specific innovation. A Foldseek search was performed to identify proteins with tertiary structure similarities, with TM-scores above 0.5 indicating conserved folds between proteins (Xu and Zhang 2010; van Kempen et al. 2024). A significant correspondence with an uncharacterized αβ-barrel domain from the crystal structure of a fungal polyglycine hydrolase was observed (TM-score = 0.73) (Dowling et al. 2023) (Fig. 2c). Authors suggested a general role of protein–protein interactions for this structural domain, though no clear functional description exists.

**Figure 2.**
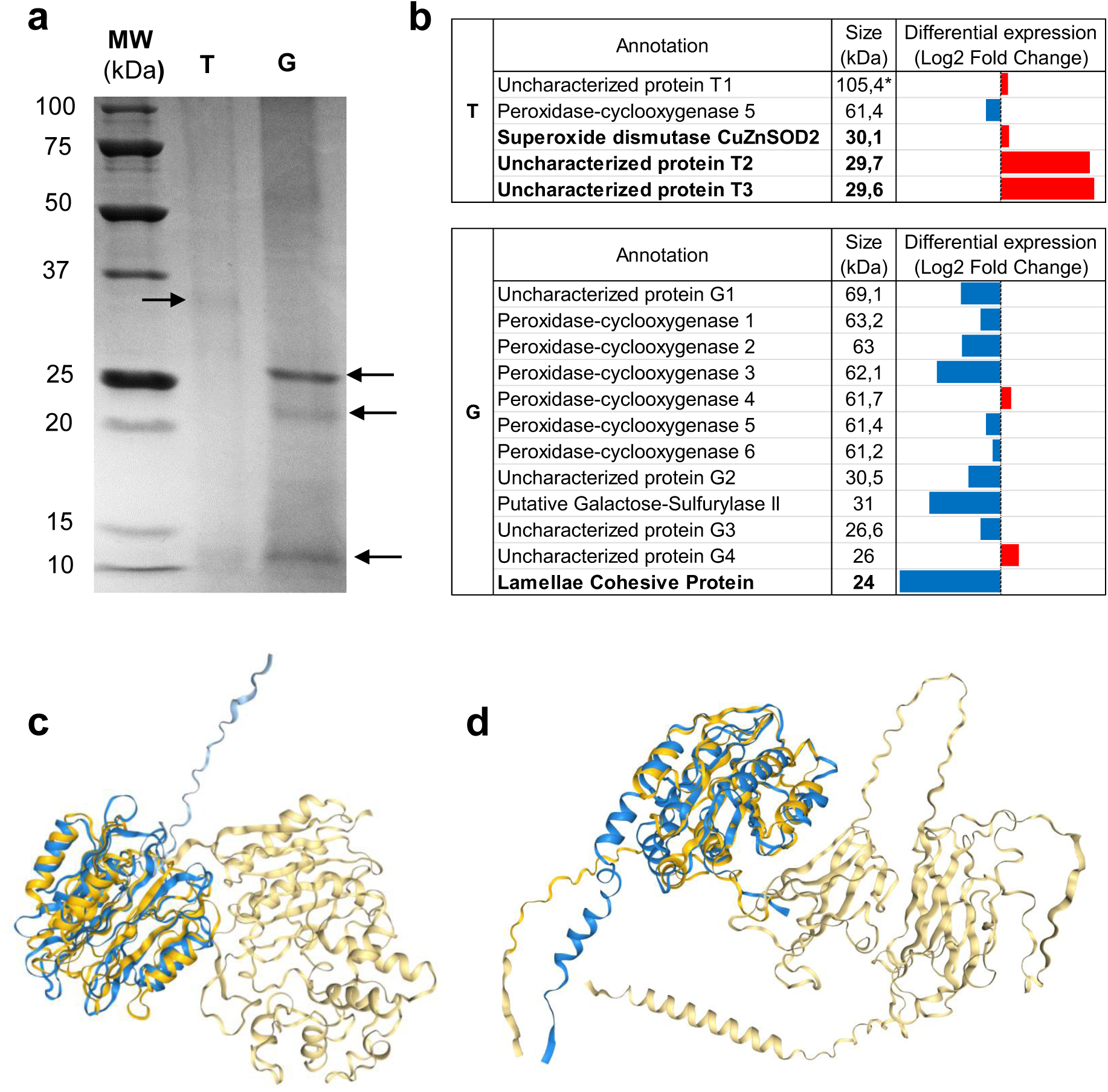
Comparative proteomic analysis of gametophyte and tetrasporophyte cuticles. **a** SDS-PAGE profiles of protein extracts from tetrasporophyte (**T**) and gametophyte (**G**) isolated cuticle samples. Black arrows indicate major bands in T and G lanes, MW: molecular weight marker. **b** LC-MS/MS analysis of trypsin digested peptide fragments of the protein extracts of T and G. Proteins identified by ≥3 peptides are shown. Column 2 refers to predicted molecular weight after signal peptide cleavage, Asterik indicates absence of signal peptide. Column 3 represents the up-regulation of the related genes in G (blue, left) or T (red, right), shown as log_2_ fold change (–11 to 11), based on data from the transcriptomic analysis of Lipinska et al. (2020). Bold names indicate candidates for the major bands. Additional details, including gene identifiers and contaminant proteins, are provided in Supplementary Table 1. **c** AlphaFold3-predicted structure of the uncharacterized protein T2 (blue) and crystal structure of a Polyglycine hydrolase from *Fusarium vanettenii* (yellow, PDB ID: 7TPU), with the overlapping of αβ-barrel domain highlighted in dark tones, as determined by Foldseek (TM-Score: 0.73, RMSD: 3.64) (https://search.foldseek.com/). **d** AlphaFold3-predicted structures of the LCP (blue) and Cuticlin-6 from *Caenorhabditis elegans* (yellow), with the overlapping of VWA highlighted in dark tones, as determined by Foldseek (TM-Score: 0.73, RMSD: 6.47).

In the gametophyte cuticle, several uncharacterized proteins have been identified, however a single protein consisting of a unique von Willebrand factor A domain (VWA) corresponds to the major band (UniProt accession: R7Q3A9), as confirmed by analysis of the excised band. VWA-containing proteins participate in a variety of functions, including in the ECM of Metazoa where VWA mediates protein–protein interactions (Whittaker and Hynes 2002). In red algae, extracellular VWA proteins have not yet been investigated, we therefore propose the name Lamellae Cohesive Protein (LCP) for the identified VWA candidate on the basis of its singular abundance in the organized multilayered cuticle and its specific expression in the gametophyte, with a log_2_ fold change of 10.7 (Lipinska et al. 2020)(Fig. 2b). The top hit from a Foldseek search was the predicted structure of Cuticlin-6, a VWA-containing protein found in the cuticle of *Caenorhabditis elegans,* showing significant similarity (TM-score = 0.73) (Fig. 2d). Cuticlin proteins are structural components that are thought to be covalently linked to form highly insoluble layers in a collagenous matrix (Fujimoto and Kanaya 1973). Other close structural similarities include important VWA-containing proteins from the Metazoan ECM, such as matrilins, collagen types VI and XXI, integrin αM, as well as the Mussel Proximal Thread Matrix Protein 1, which may oligomerized via disulfide bonds or non-covalent interactions and are involved in the binding of collagen fibrils via a metal ion dependent adhesion site (MIDAS) (Klatt et al. 2011; Emsley et al. 2000; Suhre et al. 2014; Lamande and Bateman 2018; Godwin et al. 2025). This MIDAS appears present in LCP, supported by a better result of folding prediction by addition of a divalent cation, as well as a free and exposed cysteine residues in favor of intermolecular disulfide bonds formation (Supplementary Fig. 3a, b).

The failure to identify proteins of low molecular weight, including the two other major bands of the gametophyte, may originate from an absence of trypsin cleavage sites or lack of detection by mass spectrometry. However, several proteins of higher molecular weight were positively identified, highlighting cuticle localization. These proteins include 6 enzymes of the peroxidase-cyclooxygenase superfamily (Zamocky et al. 2015), 1 putative CuZn superoxide dismutase (Hewitt and Degnan 2022), and 1 enzyme was identified as a galactose-sulfurylase (Genicot-Joncour et al. 2009) (see Supplementary Table 1 for UniProt accessions) (Fig. 2b). Peroxidase-cyclooxygenases are widespread across all domains of life, catalyzing the oxidation of diverse molecules via the reduction of H₂O₂ to water (Zamocky et al. 2015). An antimicrobial activity resulting from the production of hypohalous acids is often associated with peroxidases (Tonoyan et al. 2020). A such function supports a role in stress response or defense strategies, consistent with their accumulation in the cuticle as the first barrier with the environment. In addition, superoxide dismutases act upstream in the reaction cascade, converting O₂ to H₂O₂, and a putative member of this enzyme family was identified in the tetrasporophyte (Wolfe-Simon et al. 2005).

Another noteworthy enzyme identified in the gametophyte is a galactose-sulfurylase II family member, enzymes unique to red algae and essential for the formation of the 3,6-anhydro-D-galactose moiety found in carrageenan (Genicot-Joncour et al. 2009). This enzyme had been previously highlighted for its specific gene differential expression in the gametophyte, while many of the other members of the gene family are upregulated in the tetrasporophyte (Lipinska et al. 2020). Two galactose-2,6-sulfurylases have been biochemically characterized as key enzymes in the final step of carrageenan biosynthesis, where they catalyze the formation of 3,6-anhydro rings on sulfated galactoses (Genicot-Joncour et al. 2009). One major question regarding this reaction concerns its localization, whether carbohydrate maturation occurs in the Golgi apparatus or outside the cell (Chevenier et al. 2023). Although no other galactose-sulfurylases met the cut-off criteria in the tetrasporophyte, the experimentally identified galactose-sulfurylase II in the gametophyte cuticle supports an extracellular localization for at least this one enzyme. Nevertheless, this enzyme is relatively distant from the previously characterized enzymes of the family (Lipinska et al. 2020), and the precise specificity of this enzyme is unknown. Other proteins, whose functions are related to intracellular pathways such as gene expression or photosynthesis, were considered contaminants and excluded from further analysis (Supplementary Table 2).

### Non-protein cuticle components are mainly carbohydrates

To assess whether lipid polyester structures, such as cutin, form in the tissue, the cuticles were subjected to hydrolysis and analyzed by GC–MS. No evidence for their formation was detected (data not shown), consistent with the emergence of a specific and functional biosynthetic pathway in the divergent green lineage (Philippe et al. 2020; Kong et al. 2020). In a complementary approach to characterize other constitutive components analogous to the land plant cuticle, we analyzed soluble metabolites of the thallus surface by GC-MS, similar to waxes. Different classes of molecules were identified, including lipids and sugar derivatives, albeit present in very low amounts (Approximately 50 ng.cm^-2^) (Fig. 3a), as a comparison, the plant cuticle exhibits a wax coverage of around 1000X the metabolite content (20-200 µg.cm^-2^) (Jetter and Riederer 2016). In *C. crispus*, the most abundant molecules we detected were fatty acids and cholesterol, which are also the major lipids identified of the whole-thallus (Tasende 2000). In addition, we identified metabolites not previously described in *C. crispus*. (Z)-13-docosenamide, a compound with characterized antibacterial activity (El-Gazzar et al. 2025), as well as 3 structurally similar unknown metabolites, at highly variable levels, for which mass spectra suggest sugar derivatives with short alkylation (Supplementary Fig. 4). Biological relevance and biosynthetic origin of these molecules remain unclear.

**Figure 3.**
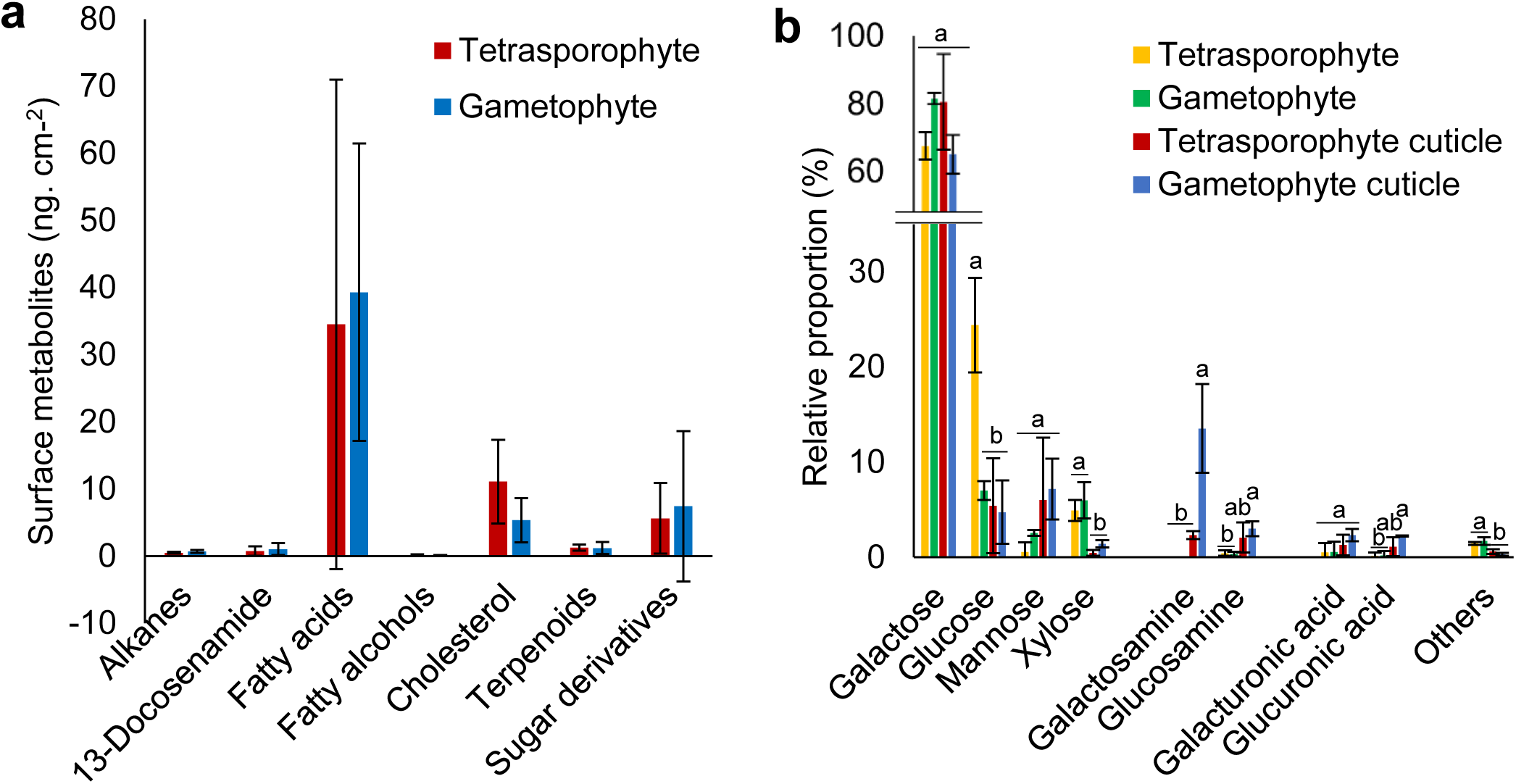
Constitutive components of the cuticle. **a** Surface metabolites from the whole thalli, extracted with nonpolar solvent. Results are the mean values ± SD of compounds detected in all samples of at least one life stage (n = 6), collected from three geographically distinct sites. No significant difference between gametophyte and tetrasporophyte (Student’s t test). All identified compounds are listed in Supplementary Table 3. **b** Sugar composition of carbohydrates from isolated cuticles and whole thalli of tetrasporophyte and gametophyte individuals. Results are the mean values ± SD from measurements of four independent cuticle isolation processes. Different letters indicate significant differences among groups (one-way ANOVA, Tukey’s HSD, p < 0.05).

To further investigate the carbohydrate constituents of the cuticle, HPAEC analysis of the hydrolyzed carbohydrate from the whole thalli and the isolated cuticles was performed, revealing a cuticle-specific sugar profile. Galactose remains the predominant sugar in the relative composition, pointing to the occurrence of galactans, very likely carrageenans, and supporting a role of the ECM polysaccharides in the assembly of the cuticle in both gametophyte and tetrasporophyte (Fig. 3b). The proportion of glucose is significantly higher in tetrasporophyte thallus, likely due to floridean starch storage (variable, not concerning), but remains similar among the other samples. However, higher proportion of mannose, lower proportion of xylose, as well as presence of amino sugars within the cuticle mark compositional differences relative to the whole thalli. In particular, galactosamine represents a substantial fraction of the sugar composition of the gametophyte cuticle, and is not detected in samples of whole thalli. In nature, galactosamine (GalN) is mainly known in its acetylated form, N-acetyl-galactosamine (GalNAc), when paired in a repetitive pattern with a uronic acid, forms the backbone of highly sulfated glycosaminoglycans (GAGs) in Metazoa. Glucuronic acid is slightly enriched in the gametophyte, suggesting an association with amino sugars. However, we currently have no information on the structure and sulfation pattern of the cuticle carbohydrates and GAG structures have never been reported in algae (Vasconcelos and Pomin 2017). Nonetheless, LCP exhibits two predicted sites of O-glycosylation that are in favor of GAG branching (Supplementary Table 1, Supplementary Fig. 3b).

Glucosamine (GlcN) is present at low levels in all samples with a slight increase in the cuticles. In the Gigartinaceae *Sarcothalia papillosa*, GlcN has been reported at significant levels in both cuticle and ECM polysaccharides (Flores et al. 1997). Moreover, in the Gigartinaceae *Mazzaella japonica*, we detected GlcN as abundant in the cuticle of both gametophyte and tetrasporophyte stages. Conversely, GalN was detected in higher proportion in the gametophyte cuticle lamellae, which shows a relatively similar SDS-PAGE protein profile to *C. crispus* and also exhibits structural coloration (Supplementary Fig. 5). Taken together, these observations indicate that a GalN content, but not GlcN, is a specific structural feature of the gametophyte cuticle, possibly associated with conserved proteins in these Gigartinaceae. Of note, using an approximately equivalent amount of isolated cuticle material, the extracted proteins were markedly less abundant in the tetrasporophyte relative to the gametophyte, conversely, the hydrolyzed sugars were significantly more abundant in tetrasporophyte samples, indicating a higher relative carbohydrate content.

### *C. crispus* LCP self-assembly exhibits affinity for carrageenans

During the development of the cuticle isolation protocol, we noted that a higher acid concentration of 70% H_2_SO_4_ led to the loss of cohesion of the lamellae, as observed by TEM, suggesting the hydrolysis of inter-lamellae components (Fig. 1g). Macromolecular composition of the isolated material was assessed by staining-based methods, revealing that the cuticle carbohydrates reactive to Toluidine Blue in the optimized protocol (35% H_2_SO_4_) were fully hydrolyzed in the altered cuticle assembly (70% H_2_SO_4_), while proteins stained with Coomassie Blue remained strongly detectable (Fig 4a, b, c, d). These results indicate a specific localization of the cuticular carbohydrates between the electron-dense layers made of proteins.

**Figure 4.**
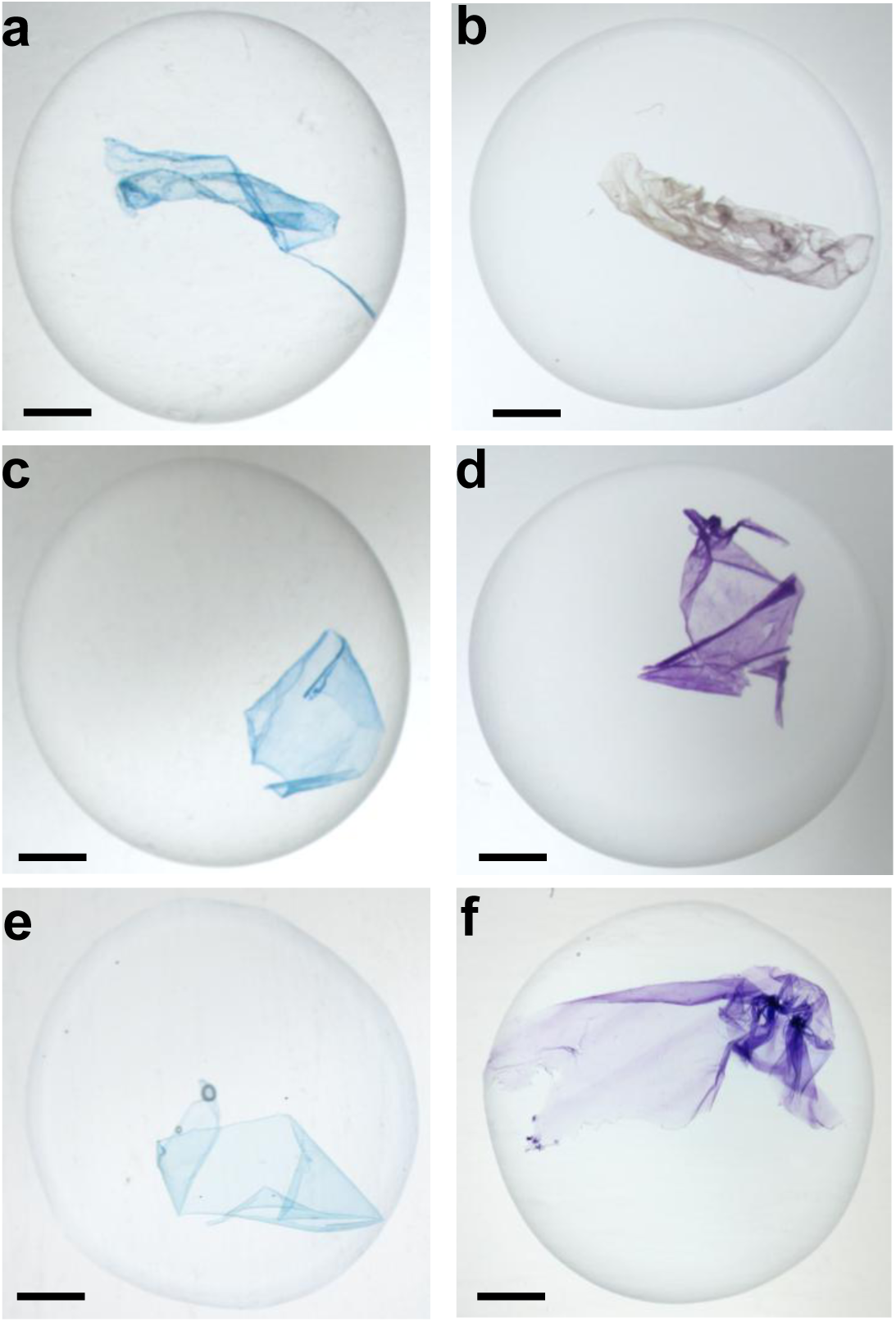
Staining assays illustrate organization of cuticle constituents. **a, b,** Isolated cuticle with an altered ultrastructure, as shown Fig. 1g. **c, d** Isolated cuticle with preserved lamellae organization, as shown Fig. 1f. **e, f** Artificial LCP film pre-incubated in carrageenan mixture **a, c, e** Coomassie blue G reveals protein content **b, d, f** Toluidine blue shows purple metachromasy when interaction with anionic polysaccharides. Scale bar = 2 mm.

To further investigate the contribution of the most abundant proteins in the cuticle assembly, a heterologous version of the LCP was produced. A denaturation step in 8M urea was necessary to successfully purify the soluble recombinant protein. Subsequently, a self-assembly of the protein occurred during the refolding step by dialysis, leading to the formation of fibril structures at the millimeter scale, resembling architecture of amyloid aggregates (Michaels et al. 2023) (Supplementary Fig. 6). Exploiting this self-assembly behavior, we produced artificial films to mimic the cuticle and test adhesion properties with carrageenans (Fig. 4e, f). We observed that the carrageenans stably coated the protein film, indicative of an interaction between the anionic sulfated polysaccharides and the positively charged surface of the LCP, which exhibits an atypically high isoelectric point of 11.7 (Fig. 4, Supplementary Fig. 3c, Supplementary Table 1). These results suggest that LCP serves a key function in connecting macromolecules within the cuticle, as well as to the ECM polysaccharides, particularly essential for anchoring the protein lamellae.

### Cuticle VWA proteins are conserved in Florideophyceae

To investigate the evolutionary relationships of LCP, we searched for homologous proteins in the NCBI database and constructed a phylogenetic tree (Fig. 5a). The LCP is also found in *M. japonica*, consistent with the highly similar protein profile of its cuticle lamellae (Supplementary Fig. 5). While only two homologs were found in *C. crispus* and *M. japonica*, numerous VWA proteins were found in species of the distant Gracilariaceae family, in which a thick, single-layer cuticle had been previously described as decklamelle (Dawes et al. 2000). Exploiting previously generated RNA-seq datasets, we obtained differential gene expression profiles for epidermal cells of *Gracilariopsis lemaneiformis* (Gracilariaceae) (Chen et al. 2022). We observed an overall up-regulation of the VWA homologs, four of the five genes exhibited a log_2_ fold change ranging from 1.8 to 3.8, consistent with a production in the epidermis of proteins involved in the cuticle assembly (Yeats et al. 2010). Chen et al. (2022) reported a higher expression of genes related to the photosynthetic metabolism in the epidermis, and lower expression for carbohydrate metabolism. Our additional examination using GO-term enrichment analysis revealed that genes only expressed in the epidermis are in most cases associated with an ECM localization, suggesting other candidates of specific roles in cuticle formation (Supplementary Fig. 7). Despite applying a relatively high detection threshold (75% coverage, 30% sequence identity), we still identified distant homologs from species with considerable evolutionary divergence. A VWA protein was found in the brown alga *Scytosiphon promiscuous* and a family of VWA-containing proteins are detected in bony fish species, all remaining functionally uncharacterized. While no VWA protein sequences were identified in other red algal families, probably due to the limited number of annotated genomes available, an analysis of transcriptomic and genomic databases revealed a widespread distribution of homologs, mainly in the large class of Florideophyceae (Fig. 5b). By contrast, in the diverging class Bangiophyceae, mostly represented by morphologically simple algae such as filaments or single cell layer sheet-like forms, no homology was found with previously reported VWA-containing proteins (Brawley et al. 2017). Nevertheless, a protein-rich cuticle had been described in the multicellular Bangiophyceae *Porphyra umbilicalis*, with a polysaccharide fraction largely consisting of mannan, thereby constituting an interesting model for further investigations (Frei and Preston 1964; Hanic and Craigie 1969).

**Figure 5.**
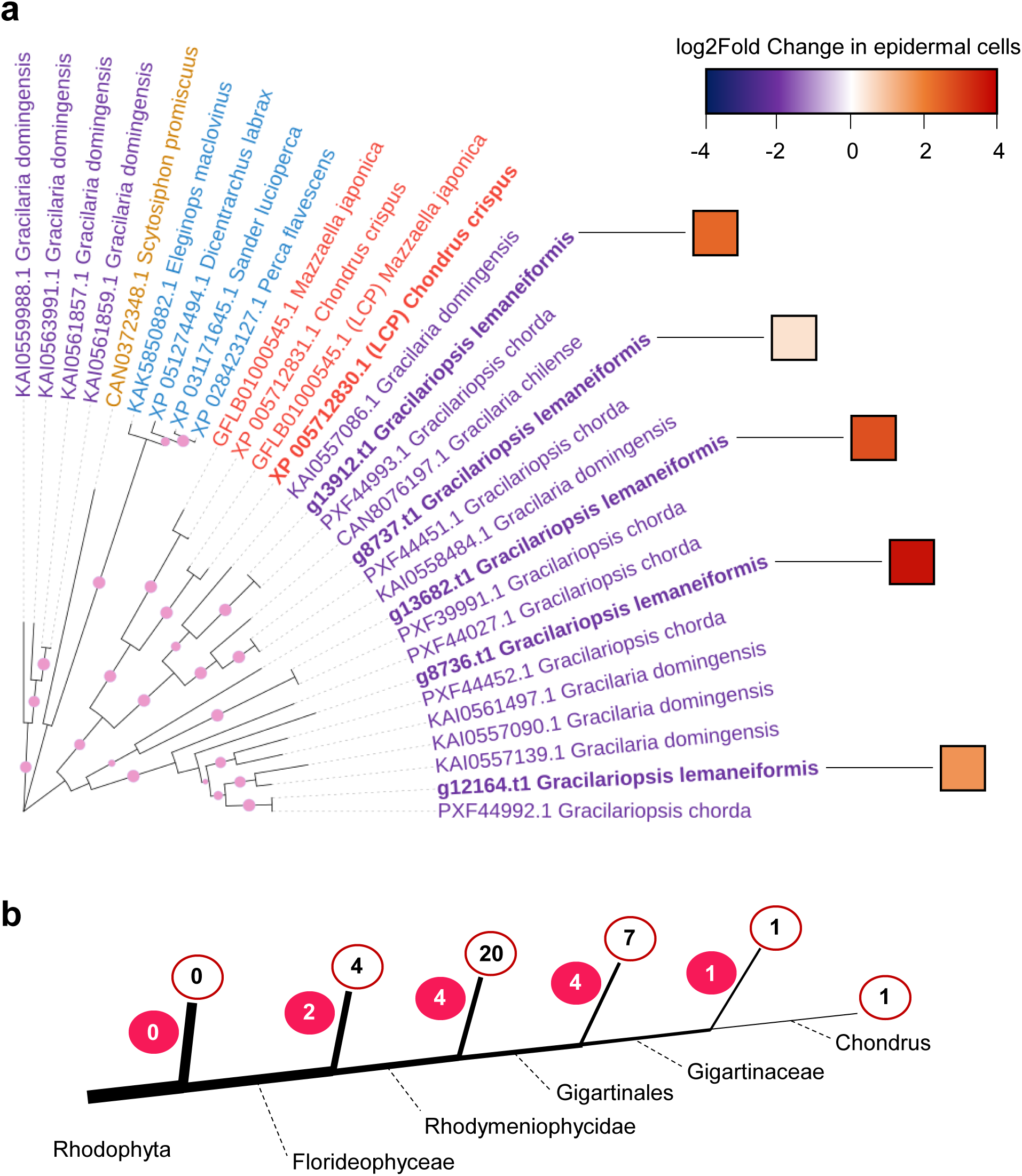
Phylogeny of LCP highlights putative orthologs. **a,** Phylogenetic tree of the LCP closest homologs identified by NCBI BLASTp, including additional related predicted proteins from *Mazaella japonica* and *Gracilariopsis lemaneiformis* genomes (Maximum likelihood analysis, bootstrap of 100 replicates). Epidermal gene expression of VWA-proteins from *G. lemaneiformis* is shown as log_2_ fold change, based on data from the transcriptomic analysis of Chen et al. (2022). Taxa in the phylogeny are represented by colored names (red, Gigartinaceae; purple, Gracilariaceae; dark yellow, brown algae; blue, bony fishes), LCP and *G. lemaneiformis* homologs are shown in bold. Branches with bootstrap values > 0.7 are marked by pink dots. **b,** Simplified phylogenetic tree of the phylum Rhodophyta, showing the number of diverging branches at each taxonomic level and the number of species in which at least one putative LCP ortholog was detected (filled red circles and open red circles, respectively). Taxonomic levels include class, subclass, order, family, and genus, as currently delineated in Algaebase (https://www.algaebase.org/). Data were obtained from NCBI tBLASTn searches of available TSA and WGS databases for Rhodophyta on NCBI. Species names are provided in Supplementary Table 4.

## Discussion

The deposition of multilayers arranged to produce structural coloration raises questions regarding how such a process is established. Understanding of the genetic pathways involved in structural coloration remains limited due to their rare distribution and predominance in non-model organisms (Airoldi et al. 2018). In plants, multilayer reflectors in the cell wall have only been reported in the epidermis of a few blue fruits (Middleton and Sinnott-Armstrong 2024). Studies have described these multilayer reflectors located beneath the plant cuticle and consist of globular lipid inclusions in the cellulose-rich cell wall (Sinnott-Armstrong et al. 2023; Sinnott-Armstrong et al. 2022). By contrast, the formation of lamellae in *C. crispus* is intrinsically linked to the cuticle organization and derived from the deposition of alternating protein and carrageenan-rich layers supported by our biochemical characterization. Protein-polysaccharide nanostructure involved in structural coloration has been described in the dermal collagen-rich ECM of some animals, leading to light interference, yet resulting of an arrangement of collagen arrays (Prum and Torres 2004; Prum and Torres 2003). This contrasts with the true multilayer thin-film produced in *C. crispus* (Fig. 1e).

Recent research supports a role for the multilayer cuticle in the protection of the photosynthetic system via the attenuation of shorter light wavelengths (Fleitas et al. 2024). It has been suggested that the specific localization of the structural coloration, at the tip of the gametophyte individuals may be associated with an underlying protection of the sexual reproductive cells (Fleitas et al. 2024; Arnould-Pétré et al. 2025). The cuticle ultrastructure has been described to vary drastically depending on the localization on the gametophyte thallus, with a noticeable absence of layers at the base (Chandler et al. 2015). In addition, an altered structure with loss of layers was reported in the middle of the thallus, suggesting a shedding induced by structural modifications or abraded surface over time. However, the growth of *C. crispus* is not restricted to the apical section (Chopin and Floc’h 1987), which suggests neo synthesis of cuticular material as the surface increases in other sections. The roles of the algal cuticle remain largely unexplored and is likely related to protective barrier function suggested by the important resistance to harsh chemical treatments (Hanic and Craigie 1969). In some brown algal species, a sustained regeneration process of the cuticle has been associated with a regulation of the epiphyte microbiome, involving the shedding of a large part of the epidermal cell wall (Halat et al. 2020; Fletcher 2024; Rousseau et al. 2025). Considering the shedding of cuticle layers in the older sections of the *C. crispus* thallus, an analogous function might be mediated by structural remodeling of the outermost part of the cuticle. Our biochemical investigations mainly focused on the highly organized lamellae of the tip section, which was better-isolated. Nonetheless, the cuticle composition may still vary in older thalli sections, and deserves future investigation.

In agreement with prior studies, no traces of cutin nor other structural lipidic polymers were found in *C. crispus,* for which enzymes of the biosynthetic pathways emerged only in the green lineage (Kong et al. 2020; Philippe et al. 2020). Moreover, presence of highly sulfated and non-acetylated polysaccharides differs from the plant cuticle, in which embedded polysaccharides have been described in favor of hydrophobic interactions (Philippe et al. 2019). Furthermore, the identified structural proteins in *C. crispus* exhibit a high surface polarity, and extremely low amount of free lipids was detected at the thallus surface, consistent with a cuticular function not related to protection against water loss. The chemical characterization of isolated *C. crispus* cuticle has revealed distinct macromolecular assemblies between the isomorphic life cycle stages. Major changes in the structure of ECM carrageenans exist between gametophytes and tetrasporophytes (McCandless et al. 1973; Ropartz et al. 2025), and their abundance extending into the cuticle are likely providing anchoring sites to specific protein constituents. Our study suggests that carrageenans have an important function in the cuticle assembly of the tetrasporophyte as galactose was a main component isolated from the cuticle, though it remains unclear if the identified tetrasporophyte αβ-barrel proteins unique to Gigartinaceae form complexes. Conversely, the identification of the abundant LCP in the gametophyte cuticle led to experiment self-assembly properties and affinity with carrageenans, which provided insights into a connective function between the protein structure and sulfated polysaccharides.

Similar to the VWA of several ECM proteins including collagen, LCP contains a characterized MIDAS that may mediate inter-protein interactions and free cysteine residue that can be involved in VWA oligomerization by disulfide bonds (Godwin et al. 2025). Interestingly, the data point to the co-occurrence of this VWA protein with GAG-like in the cuticle, as suggested by the presence of GalN along with uronic acids, thus reflecting an ECM feature of animals. The singular abundance of GalN and absence of GalNAc may indicate the activity of a putative N-deacetylase/N-sulfotransferase enzyme, which has been described in the sulfation reaction of GAGs by removing the acetyl group of a N-acetyl-glucosamine (GlcNAc) substrate (Deligny et al. 2016). The most described GalNAc-containing GAGs are likely chondroitin sulfate and dermatan sulfate, where different degrees of sulfation are associated with various functions, including structural interactions with proteins of the ECM and connective tissues in Metazoa (Mikami and Kitagawa 2013; Kjellen and Lindahl 2018). Notably, GAGs are found to interact with collagen matrix, regulating the microarchitecture of the assembly (Cortes-Medina et al. 2024). Interestingly, several genes of *C. crispus* are predicted to encode for homologs of family 7 glycosyl transferases (GT7) enzymes; some metazoan GT7 members catalyze the formation of the β-1,4 linkage between GalNAc and a D-glucuronic acid during chondroitin sulfate biosynthesis (Ficko-Blean et al. 2015). Looking at the TEM images of the gametophyte cuticle lamellae, it is tempting to draw a comparison with the GAG-rich basement membrane of the Metazoa epithelium, where narrow filaments connect the electron dense lamina to the plasmalemma (Gordon and Bernfield 1980). The additional presence of several peroxidase-cyclooxygenases within the cuticle strengthens the analogy with the basement membrane. In metazoans, enzymes of this superfamily reinforce a dense protein network by catalyzing intermolecular covalent bonds (Bhave et al. 2012); their precise function in red algae remains unknown.

It is currently unknown whether GAG-like are produced in other Rhodophyta organisms; however, genomes of Gracilariaceae and Bangyophyceae also include enzymes of the GT7 family that might be involved their biosynthesis (Brawley et al. 2017; Chen et al. 2023). An appreciable amount of mannose may also indicate a common feature with the mannan-rich cuticle of distant Bangiophyceae species (Frei and Preston 1964). In *G. lemaneiformis*, several copies of VWA protein were specifically expressed in the epidermis, supporting a shared structural function with their *C. crispus* homologs in the cuticle. Many other putative VWA protein homologs have been identified across the Florideophyceae, suggesting a conserved protein family among the taxa, but additional studies are necessary to assess their function. Although the plant cell wall contains many structural proteins, none are present in the cuticle, and VWA-containing proteins are limited to intracellular functions (Showalter 1993; Whittaker and Hynes 2002). Nonetheless, the trafficking processes of cuticular material through the ECM remain an unanswered questions for both land plants and algae (Philippe et al. 2022). The noticeable presence of electron-dense vesicle-like structures distributed throughout the epidermis may provide a basis for further exploration of cuticular material transport and deposition *in situ*. As the number of sequenced genomes in Rhodophyta continues to grow, so does the opportunity to trace the origin of structural bases for the cuticle formation (Borg et al. 2023). Nevertheless, our study offers promising candidates for genome editing in macroalgae, an approach that has yet to be consistently developed in Rhodophyta for investigating the biosynthesis and functions of the cuticle (De Saeger et al. 2024; Wang et al. 2024).

## Methods

### Cuticle isolation

Fresh, undamaged algae with sufficiently large apical tip surface were collected from the intertidal zone, cleaned of visible epiphytes, rinsed with filtered seawater, and drained on absorbent paper. Samples were immersed in 35% H_2_SO_4_ and incubated at room temperature. After 20 min with gentle, intermittent agitation, cuticle starts to peel off and can be transferred using a Pasteur pipette into 1 M Tris buffer, pH-adjusted with NaOH to achieve neutralization, for subsequent handling. The solution was replaced with distilled water at least five times to wash the samples, and gentle centrifugation between washes was applied to facilitate settling. Algae immersed in 70% H_2_SO_4_ fall apart and cuticle need to be cleared of debris after just 5min of incubation. Subsequent steps were carried out in the same manner.

### Microscopic analysis

For TEM sections, samples were fixed at 4°C for 24 hours in a fixative containing 2,5% glutaraldehyde, 0.4 M sodium cacodylate buffer (pH 7.4), and 10% NaCl. Samples were then post-fixed at 4°C for 60 minutes in a solution of 1% osmium tetroxide buffered with 0.4 M sodium cacodylate and 10% NaCl. After that samples were then dehydrated through an ethanol series (absolute anhydrous ethanol, Carlo Erba) and were embedded in Spurr resin at 60°C for two days. Sections were processed using a diamond knife on a Leica Ultracut UCT ultramicrotome, mounted on copper grids, followed by staining with uranyl acetate and lead citrate. The sections were examined and photographed using a JEOL JEM 1400 transmission electron microscope (JEOL, Tokyo, Japan) equipped with a Gatan ultrascan camera.

### Protein identification by mass spectrometry

Proteins were extracted from isolated cuticles by incubation 10 min at 100°C in Laemmli buffer containing 5% (v/v) β-mercaptoéthanol. After cooling, samples were directly loaded onto an SDS-PAGE gel for electrophoresis separation. To extract the whole cuticle proteome for proteomic analysis, electrophoresis was stopped once all proteins had entered the gel, and the upper portion was excised as a single band. Gel pieces were in gel digested following steps previously described (Yilmaz et al. 2021) with some modifications. Briefly after several washes in acetonitrile and 100 mM ammonium bicarbonate, proteins in gel piece were reduced with 65 mM DTT in 100 mM ammonium bicarbonate (15 min at 37 °C) then alkylated with 135 mM iodoacetamide in 100 mM ammonium bicarbonate (15 min at room temperature in the dark). After several washes in acetonitrile and 100 mM ammonium bicarbonate, proteins were digested with 4ng/µL of a Trypsin/LysC mix (Promega) in 25 mM ammonium bicarbonate /ProteaseMAXTM 0.01% (Promega) overnight at 37°C. Finally, peptides were collected after recovering steps in first 70% acetonitrile/0.1% formic acid and after 100% acetonitrile. Peptides were then evaporated in a vacuum centrifuge to 40 µL and purified using Phoenix spin cartridge (Preomics) according to manufacturer’s instructions.

Approximately 200 ng of the resulting peptides were analyzed by nanoLC-MS/MS using a nanoElute 2 (Bruker Daltonik GmbH, Bremen, Germany) coupled with a TimsTOF Pro mass spectrometer (Bruker) using PASEF acquisition mode as previously described (Girard et al. 2023). Ion mobility resolved mass spectra, nested ion mobility vs m/Z distributions, as well as summed fragment ion intensities were extracted from the raw data file with DataAnalysis 6.0 (Bruker Daltonik). Signal-to-noise (S/N) ratios were increased by summations of individual TIMS scans. Mobility peak positions and peak half-widths were determined based on extracted ion mobilograms (±0.05 Da) using the peak detection algorithm implemented in the DataAnalysis software. Feature detection was also performed using DataAnalysis 6.0 software and exported in .mgf format.

Peptide and protein identification were performed using the Mascot (Mascot server v2.6.2; http://www.matrixscience.com) database search engine. MS/MS spectra were queried against the complete *Chondrus Crispus* proteome UP000012073 from UniProtKB 2024_07 release, restricted to one protein sequence per gene (9598 sequences) and a common proteomic contaminant database from the Max Planck Institute of Biochemistry, Martinsried (247 sequences). Mass tolerance for MS and MS/MS was set at 15 ppm and 0.05 Da. The enzyme selectivity was set to full trypsin with one miscleavage allowed. Protein modifications were fixed carbamidomethylation of cysteines, variable oxidation of methionine. Identification results from Mascot (.dat files) were imported into the Proline Studio software (v2.2) (Bouyssié et al. 2020) which was then used for as previously described (Mear et al. 2022). Identified peptides were validated with a peptide rank of 1 and a 1% FDR based on the Benjamini-Hochberg procedure at the peptide spectrum matches level (Couté et al. 2020). Proteins identified in the negative control (side band from the same gel) were excluded from the final dataset and only proteins with 3 validated specific peptides were considered.

### Chemical analyses of surface metabolites

Six intact samples of each stage of *C. crispus* were collected from the intertidal zone at three geographically distinct sites within a 30 km area in northwestern Brittany (France). Samples were gently washed four times in 1 µm-filtered seawater in glass vials, then lyophilized. Vials containing only a small volume of filtered seawater were used as negative controls. Lyophilised *Chondrus crispus* gametophytes and tetrasporophytes were placed in a glass beaker, to which internal standards (C24 n-alkane, C22 fatty acid and C23 primary alcohol) were added. The hydrophobic surface compounds were extracted by immersing the algae in chloroform twice for 15 s. The chloroform was then evaporated at room temperature under a gentle stream of nitrogen gas, after which the extracts were silylated by adding N,O-bis(trimethylsilyl)trifluoroacetamide (BSTFA) and heating at 70 °C for 1 h. The BSTFA was subsequently evaporated, after which the extracts were redissolved in heptane/toluene (1:1 v/v) and analysed by gas chromatography coupled to mass spectrometry (GC-MS). The GC-MS analysis was performed using a Trace 1600 GC system coupled to an ISQ 7610 mass spectrometer (Thermo Scientific), with a TG-5MS capillary column (length: 60 m; diameter: 0.32 mm; film thickness: 0.25 mm). The hydrogen carrier gas flow rate was 1.2 ml min⁻¹. The oven temperature was programmed to increase from 50 °C to 210 °C at a rate of 10 °C min⁻¹, then to 250 °C at a rate of 2 °C min⁻¹, then to 330 °C at a rate of 10 °C min⁻¹, and was finally held at 330 °C for 3 min. Samples were injected in splitless mode at an injector temperature of 300 °C and the mass spectrometer was run in scan mode over 40–500 amu (electron impact ionisation), with compounds quantified based on total ion currents and internal standards.

### Chemical analyses of carbohydrates

Dried thalli, ground into a fine powder, and isolated cuticles, directly dried in the reaction tubes, were hydrolyzed with 6 M H₂SO₄ for 140 min at 37°C. The hydrolysates were then diluted to 1 M H₂SO₄ and incubated at 100°C for 120 min. Following hydrolysis, the samples were neutralized with NaOH and diluted with water. The resulting solutions were filtered prior to monosaccharide analysis. Monosaccharide composition was then analyzed using high-performance anion-exchange chromatography (HPAEC) with conductivity detection on a Dionex ICS system equipped with a CarboPac PA1 analytical column (4 × 250 mm) and a guard column of the same type (4 × 50 mm). The column was maintained at 17°C. Eluents were: A, deionized water; B, 200 mM NaOH; and C, 1 M sodium acetate. Separation was performed using a gradient program as follows: 6.5% B to 10% B over 38 min, followed by a gradient from 0 to 60% C over 7 min. This was followed by an isocratic step at 10% B and 60% C for 5 min, then column re-equilibration for 10 min at 6.5% B and 0% C. Samples were injected and eluted under these conditions, and monosaccharides were quantified by comparison with standard solutions of 10 ug/ml.

### Cloning and expression

The coding sequence of the target protein without signal peptide was codon-optimized for *Escherichia coli* expression. A polyhistidine-tag followed by an enterokinase cleavage site was designed at the N-terminus. The gene was synthesized as a custom insert and cloned into the pET-28a(+) vector (Twist Bioscience, USA). The final construct was sequence-verified by the supplier. The plasmid was transformed into chemically competent *E. coli* BL21(DE3) cells (Invitrogen) via heat shock, then cells were plated on LB agar containing 50 µg/mL kanamycin and incubated overnight at 37°C. Individual colonies were picked and screened by colony PCR to verify the presence of the insert. Verified colonies were used to inoculate 2mL pre-cultures and grown overnight at 37°C, 200 rpm, then diluted 1:100 into fresh LB medium with kanamycin for large-scale protein expression. Protein expression was induced with 0.5 mM IPTG when OD₆₀₀ reached 0.6, then cultures were incubated at 30°C for 3h with shaking at 200 rpm. Cells were harvested by centrifugation at 5,000 × g for 15 min and pellets were frozen. Cell pellets were resuspended in lysis buffer containing 20 mM Tris-HCl pH 8.0, 500 mM NaCl, and 0.1 mg/mL lysozyme, treated with DNase I to reduce viscosity, and then sonicated on ice. Pellets obtained after centrifugation at 20,000 × g were resuspended in buffer containing 50 mM Tris-HCl pH 8.0, 500 mM NaCl, 10 mM imidazole and 8M Urea, with overnight shaking at 4°C. Suspensions were clarified by centrifugation at 20,000 × g for 40 min at 4°C to remove cell debris. Supernatant was filtered through a 0.2 µm membrane filter and applied to a HisTrap FF 5mL column equilibrated with lysis buffer on an ÄKTA purifier chromatography system. The column was washed with an increase to 30 mM imidazole to remove non-specifically bound proteins, then His-tag-protein was eluted with elution buffer with a gradient of imidazole. Fractions containing the target protein were confirmed by SDS-PAGE, pooled and dialyzed against distilled water using an 8 kDa MWCO membrane to observed fibril-shaped self-assembly, concentrated and dialyzed against 20 mM Tris-HCl pH 8.0, 500 mM NaCl using 10 kDa MWCO Amicon centrifugal filters (Merck) to form protein films.

### Stain-based assays

Following dialysis by centrifugation, protein films were peeled off the membrane filter by agitation and transferred delicately into new tube avoiding contact with plastic tips. Coating by carrageenans was performed by incubation with 0.2% (w/v) of *C. crispus* purified carrageenan solution with gentle agitation for about 10 min at room temperature, then extensively washed with 20 mM Tris-HCl pH 8.0, 500 mM NaCl buffer. Isolated cuticles and protein film samples were incubated 5 min with 0.01% (w/v) toluidine blue O in buffer, or 10min with (w/v) 0.01% Coomassie brilliant blue G-250 in 3% H₃PO₄ for staining, then extensively washed with buffer. Samples were imaged under a binocular microscope.

### Phylogenetic analysis

VWA protein sequences were retrieved by BLASTp using C. crispus LCP, with the exception of VWA from *M. japonica* that were obtained from the TSA database. The sequences were aligned using M-Coffee (https://tcoffee.crg.eu/). Phylogenetic analysis was performed using maximum likelihood with 100 bootstrap replicates in MEGA11 using model parameters determined by IQ-TREE (http://iqtree.cibiv.univie.ac.at/), and the tree was visualized with iTOL (https://itol.embl.de/).

### Transcriptomic analysis

Transcriptomics data from Chen et al. (2022) were used to annotate *Gracilariopsis lemaneiformis* public genome (GCA_003346895.1). Briefly, RNA-seq reads, *de novo* transcripts (assembled with rnaSPAdes (Bushmanova et al. 2019) using both epithelial and non-epithelial data), coding sequence prediction on *de novo* transcripts (TransDecoder v5.5.0+galaxy2 - https://github.com/TransDecoder/TransDecoder) and previous CDS prediction (Sui *et al.,* personal communication) were mapped onto the *Gp. lemaneiformis* soft-masked genome (repeatmodeler and repeatmasker v2.0.3 (Flynn et al. 2020) using HISAT2 (Kim et al. 2019) and minimap2 (Li 2018). Those mapping files and a selection of public red seaweed proteomes (*Cyanidioschyzon merolae*, *Porphyridium purpureum*, *Porphyra umbilicalis*, *Chondrus crispus*, *Gracilaria domingensis*, *Gracilariopsis chorda*) were given to BRAKER3 pipeline in order to generate a new genome annotation. Newly predicted proteins were functionally annotated using both InterProScan (v5.75-106 – https://github.com/ebi-pf-team/interproscan), for function domain identification, and eggNOG-mapper v2.1.12 (Cantalapiedra et al. 2021), for protein function prediction. Protein orthology was defined based on OrthoFinder v3.1 (Emms et al. 2025) orthogroups.

Expression of newly annotated genes were quantified using featureCounts (Liao et al. 2014) and then analysed on R v4.5.2. Noised resulting from PCR amplification was removed using noisyr package v1.0 (Moutsopoulos et al. 2021) before count normalisation and differential gene expression analysis using DESeq2 v1.44 (Love et al. 2014). GO-terms enrichment was proceed using enricher tool from ClusterProfiler v4.12.6 (Wu et al. 2021).

## Acknowledgments

This study was supported by the MSCA-COFUND BIENVENUE fellowship awarded to G. Philippe from Région Bretagne and the European Union’s Horizon 2020 research and innovation program under the grant agreement No 899546, with additional funding from the Laboratory of Integrative Biology of Marine Models. We are grateful to Pr. Z. Sui for sharing genomic information of *G. lemaneiformis*, Dr. Zofia NEHR for providing PCR primers, M. JAM and A. BLIN for technical assistance, and Drs. Gurvan Michel and Philippe Potin for thoughtful discussions.

## Author Contributions

GP designed the research. GP, OG, FB, AG, DJ, SLP, EC, and LD performed experiments and analyzed the data. GP, MC, LFB and JC wrote the paper.

