## Supplementary data for "The cuticle lamellae responsible for structural coloration in red algae display common features with the metazoan extracellular matrix"

#### Supplementary data legends

**Supplementary Table 1.** Detailed table of LC-MS/MS analysis of trypsin digested peptide fragments of the protein extracts of tetrasporophyte (T) and gametophyte (G). Proteins identified by ≥3 peptides are shown. Protein molecular weights and isoelectric points calculated with (https://web.expasy.org/compute_pi/) after removal of signal peptide sequences predicted with DTU SignalP-6.0 (https://services.healthtech.dtu.dk/services/SignalP-6.0/), * indicates absence of signal peptide, O-glycosylation sites predicted with DTU NetOGlyc-4.0 (https://services.healthtech.dtu.dk/services/NetOGlyc-4.0/), subcellular localization with DTU DeepLoc-2.1 (https://services.healthtech.dtu.dk/services/DeepLoc-2.1/), and gene expression originated from the transcriptomic analysis of Lipinska et al. (2020).

**Supplementary Table 2.** List of proteins considered contaminants due to their involvement in intracellular pathways (e.g., gene expression, photosynthesis) and excluded from subsequent analyses. Proteins identified by ≥3 peptides are shown.

**Supplementary Table 3.** List of all metabolites identified on thalli surface. Values of zero are highlighted in red to indicate absence of detection; compounds not detected in any replicate were excluded from Fig. 3a and are shown in red. Asterisks indicate putative identifications based on the first NIST hit, as shown in Supplementary Fig. 4.

**Supplementary Table 4.** Species names and their phylogenetic relationships in which LCP orthologs were identified. Data were obtained from NCBI tBLASTn searches of available TSA and WGS databases for Rhodophyta on NCBI.

**Supplementary Figure 1.** Cuticle films of *C. crispus* gametophyte in suspension after chemical isolation with 35% H2SO4 (left) and 70% H2SO4 (right). Note the loss of the structural blue coloration in the right sample.

**Supplementary Figure 2.** Cuticle films of *C. crispus* gametophyte (left) and tetrasporophyte (right) in suspension after chemical isolation with 35% H2SO4. Note the thread-like aspect and absence loss of blue coloration in the right sample.

**Supplementary Figure 3.** AlphaFold predictions of the LCP without signal peptide. **a** Predicted folding in the absence or presence of divalent cations, indicated by red arrows. pTM, predicted TM-score; ipTM, interface TM-score. Higher scores in the presence of ions, highlighted in dark blue, suggest a MIDAS. **b** In the predicted structure, cysteine residues are shown in yellow, with those forming disulfide bonds highlighted as sticks; predicted O-glycosylation residues are shown in orange; beads represent $Co^{2+}$ that improves folding. **c** Surface view of the predicted structure showing electrostatic properties, with basic residues highlighted in blue and acidic residues highlighted in red. The surface shows highly basic surface patches.

**Supplementary Figure 4.** Mass spectrum of a sugar derivative compared with the top match from the NIST database. The table lists the top four hits for each of the three compounds; sugar derivatives are highlighted in yellow and red for methyl/ethyl-glucosides.

**Supplementary Figure 5.** Ultrastructure and biochemical features of *C. crispus* and *M. japonica*. **a** Gametophyte thallus of *C. crispus* (left) and *M. japonica*. (right) exhibiting structural coloration and TEM cross-sections of the multilayered cuticles at the outermost part of the ECM. **b** SDS-PAGE profiles of protein extracts from gametophyte (G) and tetrasporophyte (T) isolated cuticle of *C. crispus* (C) and *M. japonica* (M). MW: molecular weight marker. **c** Sugar compositions of carbohydrates from isolated cuticles.

**Supplementary Figure 6.** Two views of recombinant LCP showing filamentous self-assembly in suspension.

**Supplementary Figure 7.** GO-term enrichment based on genes highly upregulated in epidermal cells (Fold Change ≥ 6).

### Supplementary Table 1

| Life cycle stage | Annotation | UniProt accession | Identification score | Size (kDa) | Isoelectric Point | O-Glycosylation predicted sites | Subcellular localization prediction | Gene accession | Maximum expression in stage (TPM) | Gametophyte VS Tetrasporophyte (Log2 Fold Change) |
|---|---|---|---|---|---|---|---|---|---|---|
| Gametophyte | Uncharacterized protein G1 | R7QH99 | 130,9 | 69,1 | 8,8 | 9 | Extracellular | CHC_T00005656001 | 201 | 4 |
| | Peroxidase-cyclooxygenase 1 | R7Q2R4 | 186,8 | 63,2 | 9,7 | 19 | Extracellular | CHC_T00008635001 | 91 | 2 |
| | Peroxidase-cyclooxygenase 2 | R7QV32 | 140,0 | 63 | 8,9 | 15 | Extracellular | CHC_T00009226001 | 202 | 4 |
| | Peroxidase-cyclooxygenase 3 | R7QJ77 | 206,7 | 62,1 | 10,6 | 9 | Extracellular | CHC_T00010179001 | 1611 | 7 |
| | Peroxidase-cyclooxygenase 4 | R7QEC5 | 199,7 | 61,7 | 5,4 | 16 | Extracellular | CHC_T00010209001 | 3 | -1 |
| | Peroxidase-cyclooxygenase 5 | R7Q955 | 310,1 | 61,4 | 4,7 | 17 | Extracellular | CHC_T00008590001 | 12 | 2 |
| | Peroxidase-cyclooxygenase 6 | R7QM95 | 158,2 | 61,2 | 4,7 | 16 | Extracellular | CHC_T00010289001 | 32 | 1 |
| | Uncharacterized protein G2 | R7QNE6 | 147,3 | 30,5 | 8,57 | 37 | Extracellular | CHC_T00006690001 | 46 | 3 |
| | Putative Galactose-Sulfurylase II | R7Q2Y4 | 430,6 | 31 | 9,2 | 2 | Extracellular | CHC_T00008027001 | 1624 | 8 |
| | Uncharacterized protein G3 | R7QFM2 | 129,1 | 26,6 | 10,3 | 5 | Extracellular | CHC_T00004559001 | 784 | 2 |
| | Uncharacterized protein G4 | R7QF16 | 133,0 | 26 | 8,9 | 8 | Extracellular | CHC_T00004700001 | 93 | -2 |
| | Lamellae Cohesive Protein | R7Q3A9 | 364,6 | 24 | 11,7 | 2 | Extracellular | CHC_T00001857001 | 803 | 11 |
| Tetrasporophyte | Uncharacterized protein T1 | R7QS14 | 158,4 | 105,4* | 5,46 | 12 | Cytoplasm | CHC_T00007033001 | 94 | -1 |
| | Peroxidase-cyclooxygenase 5 | R7Q955 | 185,6 | 61,4 | 4,7 | 17 | Extracellular | CHC_T00008590001 | 4 | 2 |
| | Superoxide dismutase CuZnSOD2 | R7QRY5 | 166,3 | 30,1 | 9,4 | 10 | Extracellular | CHC_T00010182001 | 246 | -1 |
| | Uncharacterized protein T2 | R7Q339 | 231,6 | 29,7 | 9,9 | 0 | Extracellular | CHC_T00001793001 | 849 | -9 |
| | Uncharacterized protein T3 | R7Q297 | 104,2 | 29,6 | 8,7 | 0 | Extracellular | CHC_T00001406001 | 396 | -10 |

### Supplementary Table 2

| Life cycle stage | Annotation | UniProt accession | Identification score |
|---|---|---|---|
| Gametophyte | 40S ribosomal protein S6 | R7Q3M5 | 151,7 |
| | 40S ribosomal protein S16 | R7QFT3 | 145,0 |
| | Small ribosomal subunit protein uS17 | R7Q4Z1 | 117,1 |
| | Small ribosomal subunit protein uS15 | R7Q8R5 | 115,7 |
| | 40S ribosomal protein S9 | R7QCS3 | 209,4 |
| | Histone H4 | S0F310 | 199,4 |
| Tetrasporophyte | Translation elongation factor eEF1-alpha | R7Q1G0 | 114,8 |
| | F-ATPase gamma subunit | R7QTA3 | 187,2 |
| | Photosystem I reaction center subunit II | M5DBT5 | 150,2 |
| | ATP synthase CFO B chain subunit I | M5DD25 | 131,2 |
| | Prohibitin | R7QRM0 | 106,3 |
| | Histone H4 | S0F310 | 223,8 |

### Supplementary Table 3

| Class of chemicals | Compounds | Tetrasporophyte samples | | | | | | Gametophyte samples | | | | | |
|---|---|---|---|---|---|---|---|---|---|---|---|---|---|
| Linear alkanes (Alk) | 21:0 Alk | 0,03 | 0,22 | 0,16 | 0,21 | 0,04 | 0,10 | 0,12 | 0,24 | 0,14 | 0,21 | 0,08 | 0,08 |
| | 23:0 Alk | 0,00 | 0,00 | 0,00 | 0,00 | 0,00 | 0,00 | 0,00 | 0,11 | 0,31 | 0,69 | 0,00 | 0,00 |
| | 25:0 Alk | 0,00 | 0,00 | 0,00 | 0,00 | 0,00 | 0,31 | 0,00 | 0,29 | 0,68 | 0,27 | 0,00 | 0,00 |
| | 27:0 Alk | 0,00 | 0,00 | 0,00 | 0,43 | 0,00 | 0,55 | 0,00 | 0,00 | 0,69 | 0,66 | 0,00 | 0,00 |
| | 28:0 Alk | 0,27 | 0,34 | 0,07 | 0,32 | 0,06 | 0,48 | 0,20 | 0,35 | 0,82 | 0,41 | 0,00 | 0,00 |
| | 29:0 Alk | 0,00 | 0,00 | 0,00 | 0,00 | 0,00 | 0,39 | 0,00 | 0,17 | 0,51 | 0,14 | 0,00 | 0,07 |
| | 30:0 Alk | 0,00 | 0,00 | 0,00 | 0,00 | 0,00 | 0,29 | 0,00 | 0,00 | 0,39 | 0,16 | 0,00 | 0,06 |
| | 19:1 Alk | 0,00 | 0,00 | 0,00 | 0,00 | 0,00 | 0,00 | 0,00 | 0,16 | 0,66 | 0,24 | 0,00 | 0,09 |
| | 20:1 Alk | 0,00 | 0,00 | 0,00 | 0,00 | 0,00 | 0,00 | 0,00 | 0,55 | 1,10 | 0,54 | 0,00 | 0,24 |
| Fatty amides | 13-Docosenamide | 0,77 | 1,08 | 0,17 | 0,22 | 0,23 | 1,35 | 0,12 | 1,59 | 0,99 | 2,54 | 0,48 | 0,49 |
| Fatty acids | 14:0 FA | 9,32 | 22,75 | 1,21 | 0,85 | 6,92 | 1,61 | 4,85 | 17,77 | 9,54 | 3,39 | 4,01 | 2,40 |
| | 16:0 FA | 28,74 | 72,76 | 10,73 | 2,06 | 22,65 | 13,79 | 21,78 | 25,15 | 51,08 | 22,04 | 20,05 | 15,51 |
| | 18:1 (d9c) FA | 0,70 | 5,82 | 0,49 | 0,17 | 0,17 | 0,17 | 1,58 | 4,72 | 10,10 | 12,95 | 1,59 | 2,95 |
| | 18:1 (d9t) FA | 0,78 | 7,95 | 0,00 | 0,00 | 0,95 | 0,00 | 7,69 | 0,77 | 1,64 | 0,00 | 0,01 | 0,05 |
| | 18:0 FA | 0,40 | 4,32 | 0,00 | 0,00 | 0,00 | 0,00 | 0,94 | 0,00 | 8,78 | 2,73 | 1,55 | 0,00 |
| | 20:5 (5,8,11,14,17) FA | 4,27 | 10,75 | 2,61 | 2,88 | 0,48 | 2,05 | 3,23 | 8,93 | 7,18 | 5,60 | 2,52 | 1,31 |
| Monoacylglycerols | 16:1 2-MAG | 1,32 | 0,67 | 0,00 | 1,60 | 1,23 | 1,48 | 0,00 | 1,34 | 0,00 | 0,00 | 1,23 | 1,90 |
| | 16:1 1-MAG | 1,07 | 0,85 | 0,00 | 0,00 | 0,00 | 1,37 | 0,00 | 0,25 | 1,01 | 0,00 | 0,00 | 0,31 |
| | 16:0 1-MAG | 1,19 | 2,54 | 0,00 | 0,67 | 0,17 | 0,00 | 0,00 | 2,53 | 5,03 | 0,00 | 0,00 | 0,28 |
| | 18:0 1-MAG | 0,33 | 0,02 | 0,00 | 0,00 | 0,00 | 0,00 | 0,00 | 0,00 | 0,00 | 0,00 | 0,00 | 0,00 |
| Primary alcohols | 18:0 PA | 0,00 | 0,17 | 0,00 | 0,00 | 0,00 | 0,00 | 0,00 | 0,00 | 0,25 | 0,00 | 0,00 | 0,00 |
| | 22:0 PA | 0,00 | 0,24 | 0,20 | 0,14 | 0,09 | 0,28 | 0,09 | 0,07 | 0,19 | 0,03 | 0,07 | 0,07 |
| Sugar derivatives | Heptyl 1-thio-β-D-glucopyranoside* | 1,27 | 2,75 | 1,30 | 3,95 | 0,11 | 0,25 | 0,40 | 0,53 | 2,60 | 8,56 | 0,61 | 0,77 |
| | 1-Ethylhexyl β-D-glucopyranoside* | 2,05 | 2,79 | 1,18 | 6,81 | 0,08 | 0,16 | 0,49 | 0,63 | 2,53 | 15,02 | 1,02 | 1,00 |
| | (Z)-3-Hexenyl β-glucopyranoside* | 0,84 | 2,02 | 1,06 | 4,32 | 0,04 | 0,05 | 0,09 | 0,29 | 1,58 | 6,22 | 0,36 | 0,60 |
| Sterols | Cholesterol | 15,49 | 21,76 | 8,32 | 6,07 | 8,47 | 5,92 | 1,34 | 9,23 | 9,41 | 4,91 | 3,44 | 3,72 |
| Terpenoids and derivatives | Neophytadiene | 0,94 | 1,18 | 0,10 | 0,07 | 0,10 | 0,18 | 0,05 | 0,62 | 0,31 | 0,13 | 0,09 | 0,10 |
| | Phytol | 0,65 | 0,74 | 0,18 | 0,05 | 0,16 | 0,37 | 0,30 | 2,25 | 0,98 | 0,40 | 0,16 | 0,27 |
| | α-Tocopherol | 0,11 | 0,28 | 0,00 | 0,00 | 0,00 | 0,00 | 0,00 | 0,00 | 0,33 | 0,00 | 0,00 | 0,00 |

### Supplementary Table 4

| Class | Subclass | Order | Family | Species |
|---|---|---|---|---|
| **Florideophyceae** | **Rhodymeniophycidae** | **Gracilariales** | Gracilariaceae | *Gracilaria caudata, Gracilaria vermiculophylla, Gracilaria gracilis, Gracilaria domingensis, Gracilaria salicornia, Gracilaria chilensis, Gracilariopsis lemaneiformis, Gracilariopsis chorda* |
| | | **Gigartinales** | Gigartinaceae | *Chondrus crispus* , *Mazzaella japonica* |
| | | | Endocladiaceae | *Endocladia muricata* , *Gloiopeltis furcata* |
| | | | Dumontiaceae | *Dumontia simplex* |
| | | | Solieriaceae | *Kappaphycus alvarezii* , *Kappaphycus striatus* , *Eucheuma denticulatum* |
| | | | Petrocelidaceae | *Mastocarpus papillatus* |
| | | **Bonnemaisoniales** | Bonnemaisoniaceae | *Asparagopsis taxiformis* , *Bonnemaisonia californica* |
| | | | Delesseriaceae | *Delisea pulchra* |
| | | **Ceramiales** | Rhodomelaceae | *Chondria armata* , *Digenea simplex* , *Bostrychia moritziana* , *Pterothamnion heteromorphum* |
| | | | Ceramiaceae | *Herpochondria borealis* |
| | | **Halymeniales** | Halymeniaceae | *Grateloupia filicina* , *Grateloupia livida* , *Grateloupia catenata* , *Grateloupia turuturu* |
| | **Corallinophycidae** | **Sporolithales** | Sporolithaceae | *Sporolithon* sp. |
| | | **Corallinales** | Corallinaceae | *Amphiroa fragilissima* |
| | | | Hapalidiaceae | *Lithothamnion corallioides* |
| | **Nemaliophycidae** | **Palmariales** | Rhodophysemataceae | *Rhodonematella subimmersa* |
| **Bangiophyceae** | | | | |

Supplementary Figure 1

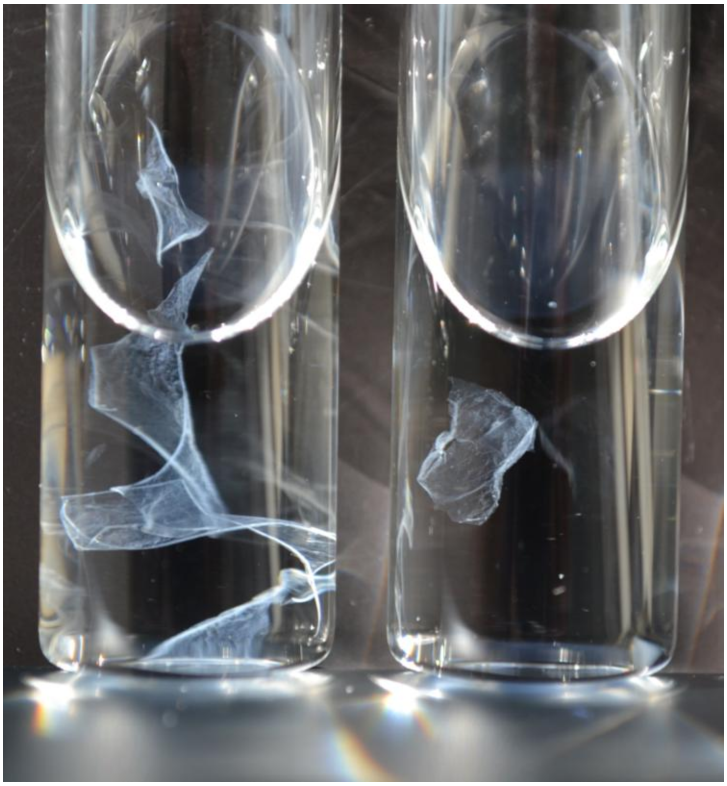

Supplemental Figure 2

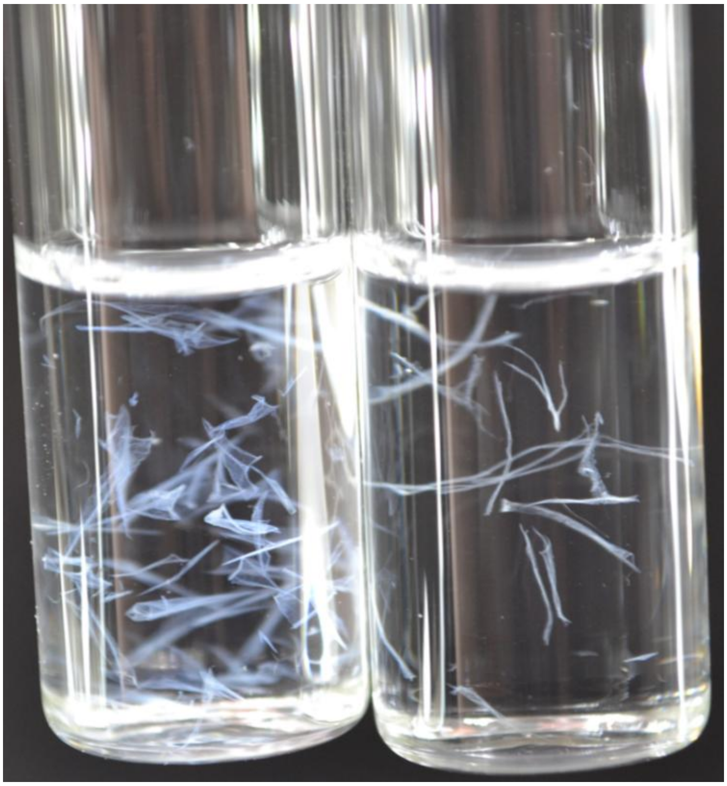

### Supplementary Figure 3

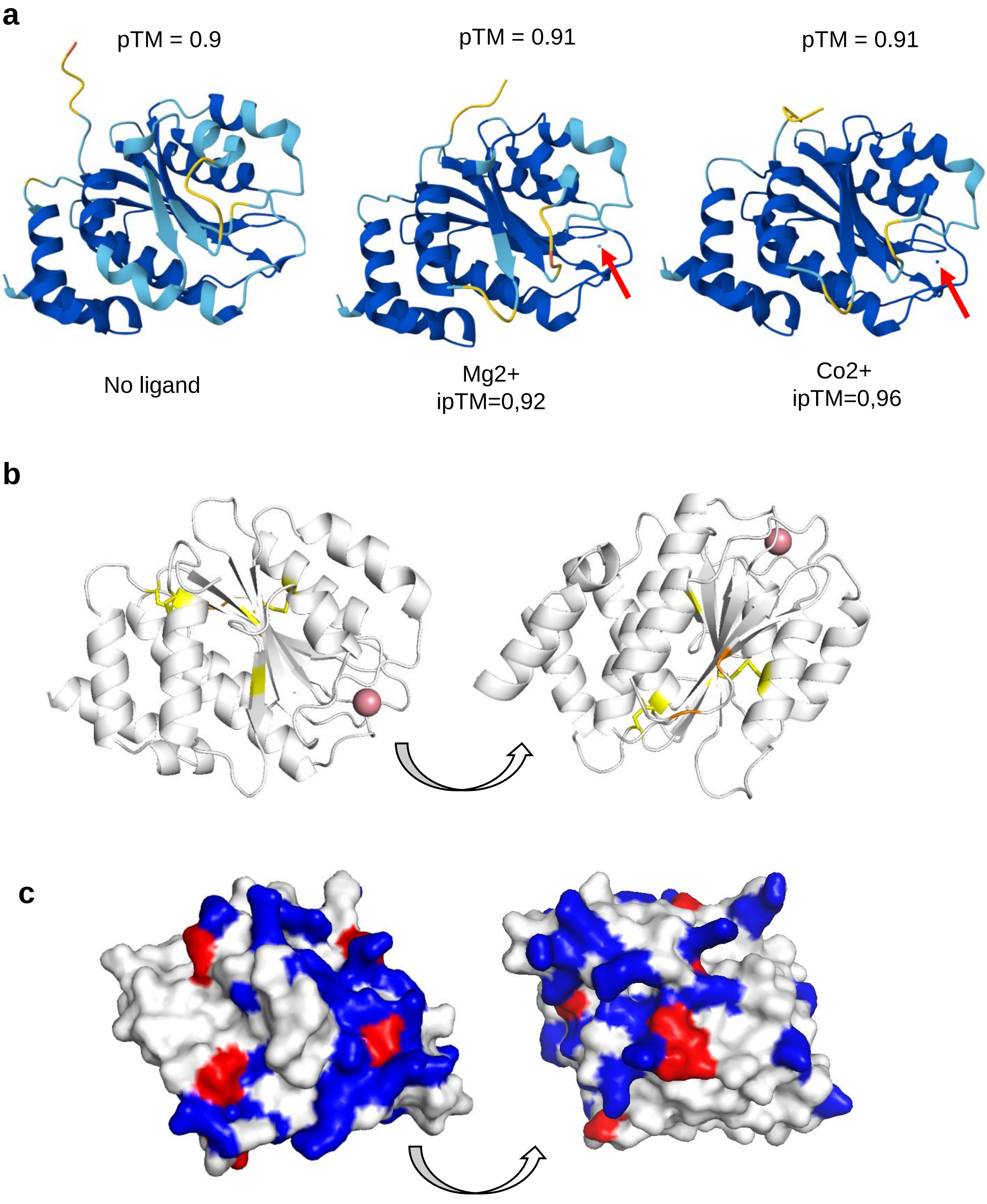


Comment: should add surface hydrophobicity?

### Supplementary Figure 4

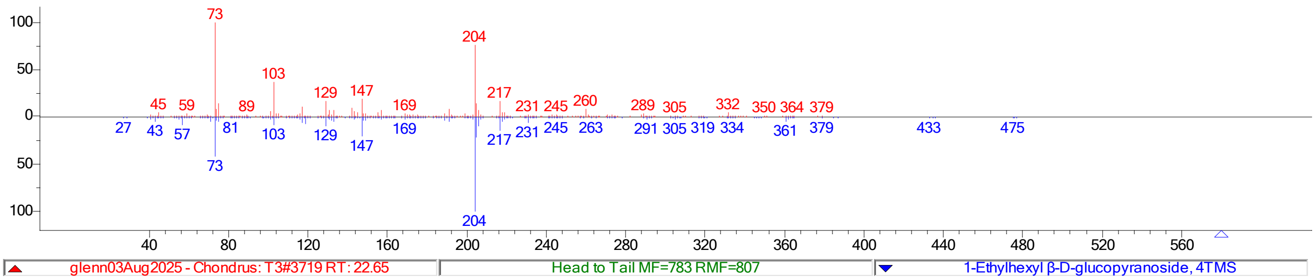


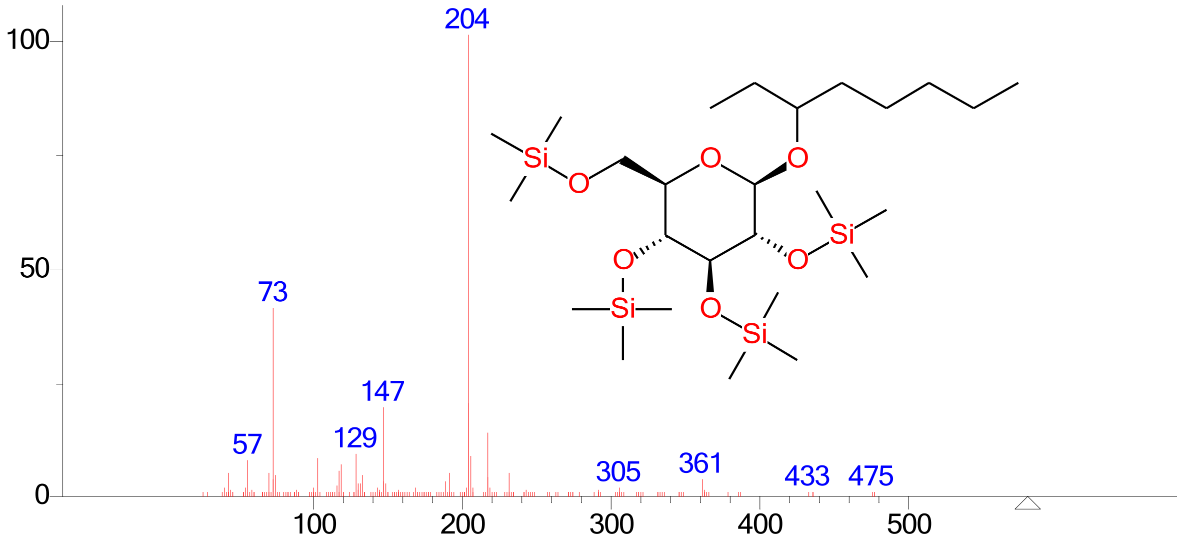


(mainlib) 1-Ethylhexyl β-D-glucopyranoside, 4TMS

| Hit number | Compound 1 | Compound 2 | Compound 3 |
|---|---|---|---|
| 1 | Heptyl 1-thio-β-D-glucopyranoside | 1-Ethylhexyl β-D-glucopyranoside | (Z)-3-Hexenyl β-glucopyranoside |
| 2 | Methyl α-Arabinofuranoside | B-D-Lactose | Methyl α-D-glucofuranoside |
| 3 | Deoxyribopyranose | Ethyl α-D-glucopyranoside | Methyl α-Arabinofuranoside |
| 4 | (Z)-3-Hexenyl β-glucopyranoside | β-D-Galactopyranoside, methyl | β-D-Tagatopyranose |

### Supplementary Figure 5

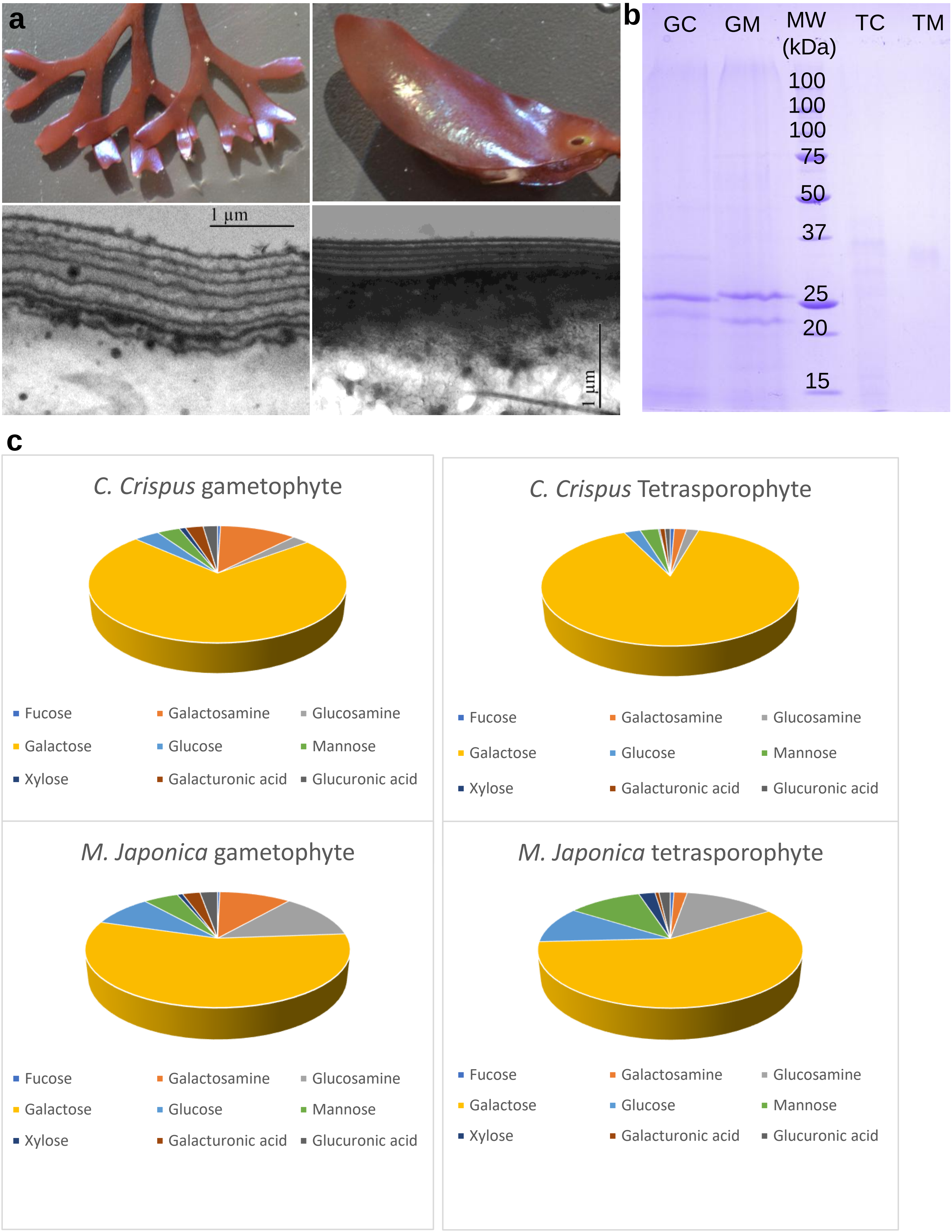

Supplementary Figure 6

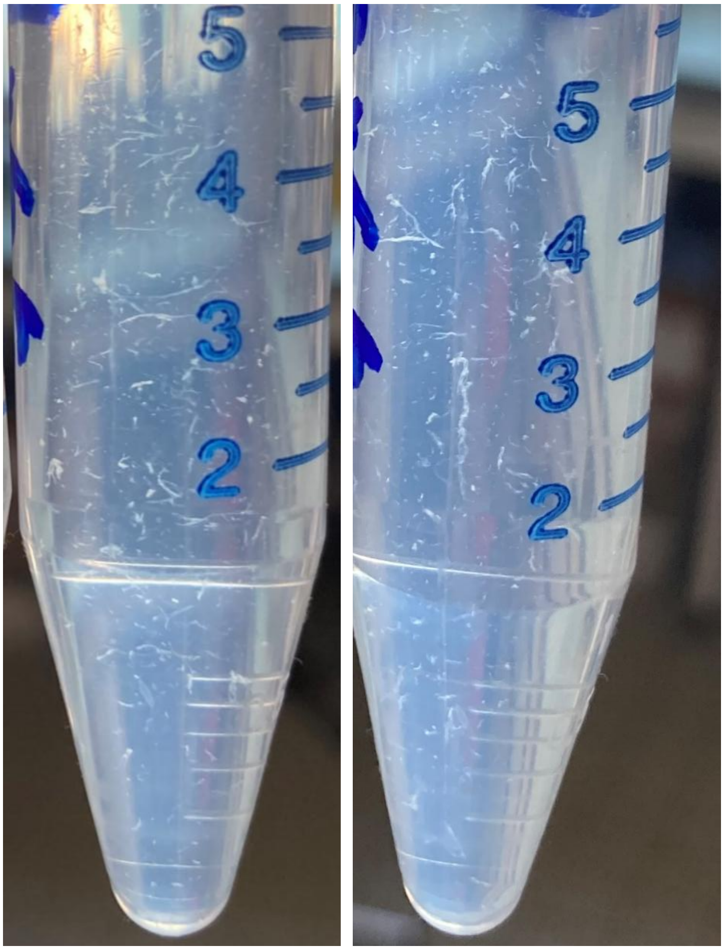

### Supplementary Figure 7

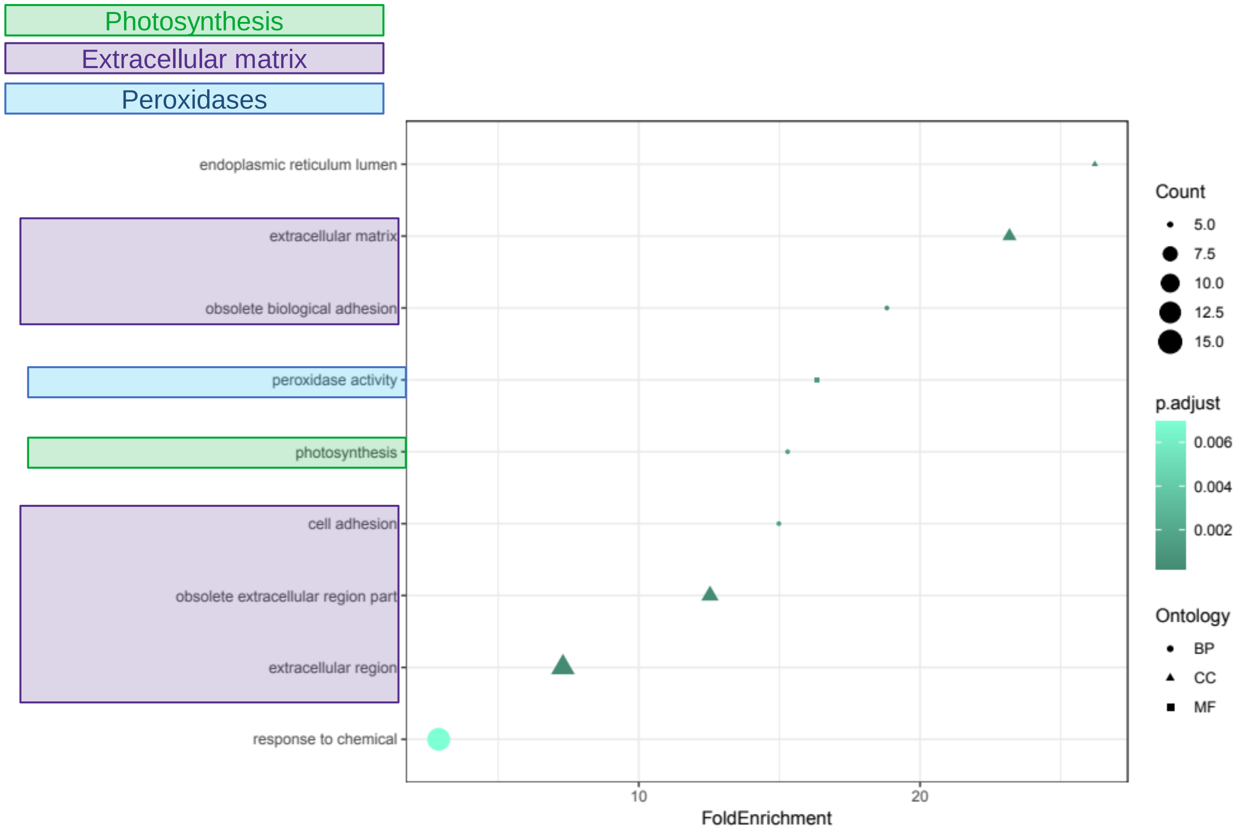
